# Co-Hyperpolarized Dehydroascorbate and Pyruvate MRI Predicts Treatment Response in Glioblastoma

**DOI:** 10.1101/2025.11.11.687898

**Authors:** Elizabeth Coffee, Paola Porcari, Saket Patel, Grace Figlioli, Marjan Berishaj, Ingo K. Mellinghoff, Kayvan R. Keshari

**Author notes:** **Corresponding Author:** Kayvan R. Keshari, PhD. 417 E 68^th^ St., New York, NY 10065. These authors contributed equally to this work (Paola Porcari, Saket Patel).

## Abstract

**Purpose:** Early noninvasive assessments of treatment response are desperately needed to improve outcomes in glioblastoma (GBM). Molecular imaging techniques that measure glycolytic metabolism are being increasingly studied, but limitations such as variable substrate delivery present significant barriers to clinical interpretation. To develop more robust translational imaging biomarkers, we propose utilizing the interrogation of oxidative stress, a critical component of tumor metabolism for which no method of clinical measurement currently exists. This study investigates the simultaneous measure of oxidative stress and glycolytic flux using co-hyperpolarized [1-^13^C] dehydroascorbate and [1-^13^C] pyruvate (HP DHA/PA) as a predictor of treatment response in GBM.

**Experimental Design:** To establish a model that exhibits known metabolic responses to oxidative stress, we characterize radiation induced metabolic reprogramming in four human GBM lines (U87, U251, A172, T98) *in vitro*. We extend this in vivo and establish radiosensitive and radioresistant orthotopic xenograft models to investigate HP DHA/PA magnetic resonance imaging as a predictor of treatment response.

**Results:** *In vitro* analyses revealed that radiation upregulates the pentose phosphate pathway and response is augmented by glutathione depletion. *In vivo* metabolomic profiling identified preferential nucleotide metabolism pathways in each tumor type. HP DHA/PA imaging revealed that DHA perfusion was not impacted by blood-brain-barrier integrity and detected reductions in DHA-to-vitamin C and pyruvate-to-lactate conversion in treatment-sensitive tumors, reflecting diminished reductive capacity following radiation.

**Conclusions:** These findings demonstrate successful prediction of radiosensitivity in GBM utilizing measurement of oxidative stress and establish HP DHA/PA imaging as an innovative method to address existing clinical limitations in treatment response assessment.

## Introduction

Glioblastoma is most common primary malignant brain tumor in adults, and the most resistant to treatment. Despite aggressive multimodality treatment, median overall survival remains effectively unchanged since the implementation of standard of care chemoradiation after maximal safe resection two decades ago.^1^ A major limitation to the advancement of the field is our inability to assess treatment response in real time. Current response assessments are limited to visualization of anatomic stability on contrasted magnetic resonance imaging (MRI), a process that requires months of surveillance. MRI changes from treatment effects (pseudoprogression) or antiangiogenic therapy (pseudoresponse) complicate interpretation of these assessments. There is, therefore, a distinct unmet clinical need for improved biomarkers for early assessment of treatment response in glioblastoma.

One avenue to assess treatment response is to quantify oxidative stress, a hallmark effect of ionizing radiation treatment. Conventional external beam radiation is standard of care in the upfront treatment of glioblastoma, and advances in radiopharmaceuticals have accelerated the need for spatially resolved measurements of treatment efficacy.^2^ Beyond radiation, sensitivity to the standard-of-care treatment temozolomide has been linked to increased reactive oxygen species (ROS)^3^ and numerous compounds that impact cellular redox balance in GBM are under investigation.^4^ Despite the key role of oxidative stress in human disease, there are no clinically available methods to noninvasively quantify this component of tumor metabolism.

Molecular imaging is a minimally invasive method for probing biological processes at the cellular and molecular level and is being increasingly utilized to address the need for earlier read outs of treatment response. Hyperpolarized (HP) MRI is one such molecular imaging method that increases the lifespan and visibility of endogenous substrates, and when paired with carbon labeling magnetic resonance spectroscopy, permits the real-time quantification of metabolic reactions in vivo.^5^ Hyperpolarized [1-^13^C] pyruvate is the most commonly utilized HP MRI imaging probe with > 60 published human studies.^6^ However, the biologic interpretation of pyruvate’s conversion to lactate is confounded by perfusion driven substrate delivery, transporter dependence, and differentiation of intra vs extracellular lactate pools. Beyond pyruvate, HP [1- ^13^C] dehydroascorbate has been utilized as a redox sensor in preclinical studies to quantify oxidative stress in a variety of disease states.^7-15^

To address the need for more robust measurements of tumor metabolism that improve upon current clinical methods, we present the use of co-hyperpolarized dehydroascorbate and pyruvate (HP DHA/PA) to predict treatment response to radiation in orthotopic models of glioblastoma.

## Methods

### Cell culture

Human U87-MG (RRID:CVCL_0022), A172 (RRID:CVCL_0131), T98 (RRID:CVCL_B368) human male glioblastoma cell lines (American Type Culture Collection, Manassas, VA, United States) and U251 glioblastoma cell line (gift from the laboratory of Ingo Mellinghoff, MD and authenticated via STR analysis October 2025) (RRID:CVCL_0021), were used. Cells were cultured in Dulbecco’s modified Eagle’s medium without glucose, supplemented with 10% fetal bovine serum (Gibco BRL, Thermo Fisher Scientific), penicillin (50 U/ml) and streptomycin (250 μg/ml; Gibco), and 5.5 mM glucose (unless otherwise noted) in a humidified 5% CO_2_/95% air at 37°C. Cells were tested for mycoplasma using MycoAlert® Mycoplasma Detection Kit Unless (Lonza LT07-318) every month and used only at low passage (<10).

### Radiation response in vitro

Radiation was administered via X-ray irradiator XRAD-320 (Precision X-Ray) at a rate 117 cGy/min. Viability assays were performed by counting viable cells 72 hours after treatment using trypan blue staining. Cells treated with L-Buthionine-(S,R)-Sulfoximine (Cayman Chemical, 14484) were treated with 500uM for 18 hours preceding radiation treatment unless otherwise noted. Proliferation assays were performed on IncuCyte SX5 (Sartorious) and confluence was normalized to starting confluence per well before normalizing to untreated control. All experiments were performed in biologic triplicates.

### Colony Forming Assay (CFA)

CFA were performed by plating 1000 cells/well in a 6-well plate from serial dilution of a single concentrated cell suspension. Wells were treated with 100uM BSO for 18 hours, 10 Gy radiation, or both and maintained in high glucose DMEM (25 mM). After 14 days, cells were fixed with 4% paraformaldehyde and stained with 0.1% crystal violet. Colonies containing > 50 cells were counted manually.

### Carbon Tracing

Cells were treated with 5 mM [1,2-13C] glucose (Sigma 453188) in media without glucose and media collected after 4 hours. Media samples were prepared with a standard media buffer solution, 5mM [1-^13^C] urea for concentration quantification, and 2uL of gavodvist in 500 uL solution and run in a ^13^C NMR at 600 MHz (Avance III). NMR spectroscopy data were processed using the Mnova Software Suite (version 14.3.2, Santiago de Compostela, Spain). Lactate signals centered at 20.1 ppm were normalized to 1-^13^C urea at 160ppm, and line fitting within Mnova used to calculate integral under each peak.

### Metabolomics

GBM cells (triplicate for each condition) or tumor samples (n=3 mice for each of 4 conditions) were mixed or homogenized in cold (−80 °C) 80% methanol. After centrifuging, samples were lyophilized by speed vac. Dried pellets were resuspended in 1:1 methanol:H2O before LC-MS/MS analysis and normalized volumes used. Metabolites were separated by Hydrophilic interaction chromatography (HILIC) and detected by high-resolution mass spectroscopy (5ppm mass accuracy). Samples were run in triplicate on a Vanquish Horizon / Q-Exactive HF Orbitrap LC-MS system. Untargeted metabolite profiling of cell and tissue extracts was performed first using definitive identification based on the mass and retention time of authentic chemical standards, followed by putatively annotated compounds by searching HMDB database (RRID:SCR_007712) using accurate mass. MetaboAnalyst 6.0 (RRID:SCR_015539) was used for data processing. Raw intensities were normalized to median and log2 transformed. Pathway enrichment analysis was conducted first by selecting metabolites using a threshold of p<0.05 and Fold Change > 2 and Q-stat based on established methods^16^, run against a library of SMPDB based on normal human metabolic pathways. Pathways enriched with p> 0.1 were included. Individual metabolite pool values were compared using normalized data before log transformation. Volcano plots for tumor samples were generated using thresholds of p<0.1 and FC >1.5 (<0.66) in tumor samples due to a wide variability in sample intensities after normalization. Raw data is included as a supplementary file.

### Orthotopic Xenografts

All animal studies were done in accordance with IACUC approved protocol. Human glioblastoma cells were orthotopically implanted in athymic Foxn1nu mice (Jackson Laboratories, RRID:IMSR_JAX:007850). Anesthetized mice were placed into a stereotaxic frame (Stoelting Co., Wood Dale, IL, United States), and 3 × 10^5^ cells in suspension were injected in a volume of 2 μl of phosphate-buffered saline via a stereotaxic injector at a flow rate of 2 μl/min into the right striatum of the mice (coordinates from bregma: 0.0-mm posterior, 2.5-mm lateral, and 3.0-mm ventral).

### Spectroscopic Imaging

Four groups of athymic nude mice (Jackson Laboratories, RRID:IMSR_JAX:007850) were imaged (n=3 of each U251 and U251 RT and U87, n=4 of U87 RT). Spectroscopic imaging experiments were performed on a Bruker BioSpec 3.0-T/18-cm horizontal bore system (Bruker BioSpin, Billerica, MA). A dual-tuned 1H/^13^C transmit/receive radio frequency volume coil (Bruker BioSpin MRI GmbH, Ettlingen, Germany) was used. T2-weighted images of mouse brain were acquired in the coronal orientation as an anatomical reference. Static magnetic field homogeneity was optimized using a localized field-map shimming routine (ParaVision 360, v2.0 pl 1, Bruker BioSpin, RRID:SCR_025295)

After initial imaging sequences, anesthesia was held before injection with the HP solution. Each mouse underwent the ^13^C hyperpolarized MRI scan 15 s after injecting 150 μl of co-hyperpolarized 40 mM [1-^13^C]DHA, 90 mM [1-^13^C]pyruvate over 10 seconds. 2D multivoxel CSI sequence was acquired coronally (TR/TE_eff_ = 141.4/5 ms, field of view = 32 × 32 mm^2^, flip angle of 20°, in-plane resolution of 2.667 × 2.667 mm^2^, 5 mm slice thickness centered over the brain tumor). The ^13^C CSI sequence was repeated on a cylindrical ^13^C urea phantom (6M) for signal normalization.

### Hyperpolarized Data analysis

Spectroscopic data were preprocessed through a custom Matlab script (Matlab R2021b, The MathWorks, RRID:SCR_001622) applying 10 Hz apodization, zero-filling interpolation (ZFP), baseline correction. Processed spectra were then overlayed on anatomic images in Spectroscopic Imaging Visualization and Computing (SIVIC 0.9; RRID:SCR_027875) software package.^17^ Integration regions selected for each of: Pyruvate, DHA, Vitamin C, Pyruvate Hydrate, and Lactate over tumor and contralateral (CL) brain ROIs. Mean values of each metabolite were normalized to the 6 M ^13^C urea phantom, measured polarization and concentrations of injected substrates.

### Tissue extraction

Mice were sacrificed utilizing isoflurane overdose in accordance with IACUC approved animal protocol. Mouse brains were rapidly extracted under cold condition and were snap frozen either whole or after tumor dissection in liquid nitrogen. Whole brain samples for tissue staining were sectioned at -20C in 10 μm slices. Immunohistochemistry with Ki-67 Rabbit mAb (Cell Signaling Technology Cat# 9027, RRID:AB_2636984) at 2 μL/ml was performed. Immunofluorescence was performed with with Anti-gamma H2A.X (phospho S139) antibody (Abcam Cat# ab11174, RRID:AB_297813) and Alexa Fluor 594.

### Statistical Analysis

Data analyses were performed using GraphPad Prism (RRID:SCR_002798). A two-tailed paired or unpaired Student’s t test was used to determine significance when two conditions were compared and is noted in the text. For experiments with more than two conditions a one or two-way ANOVA was used. For survival data a log rank test was used. In all cases values of *p* < 0.05 were considered significant. *p* < 0.05; **, *p* < 0.01; ***, *p* < 0.001; ****, *p* < 0.0001. Data are shown as mean ± SEM (standard error of the mean).

## I. Results

### A. Glutathione Depletion Increases Glioblastoma Sensitivity to Radiation

To identify differences in the response to oxidative stress, we examined four immortalized glioblastoma cell lines. First, we examined viability at 72 hours after treatment with ascending doses of ionizing radiation: 0 Gy, 2 Gy, 4 Gy, and 10 Gy (Fig 1A) to estimate an IC50 for each cell line. T98s exhibited the highest resistance to treatment (IC50=7.2 Gy, R^2^=0.75). At the cellular level, the equilibrium between reductive and oxidative electron transfer reactions is critical for maintaining cellular functions. Reduced glutathione (GSH) is a the most abundant neutralizer of reactive oxygen species (ROS) and its’ balance with oxidized glutathione (GSSG) is a marker of cellular oxidative stress.^18^ Radiation causes dysregulation of cellular redox status and oxidative stress via the formation of free radicals which impacts downstream signaling pathways, leading to changes in cell growth, differentiation, and cell death.^19^ We therefore examined the effects of glutathione depletion on the sensitivity to radiation in glioblastoma cell lines (Fig 1B-E). Buthionine sulfoximine (BSO) is a potent irreversible inhibitor the rate-limiting enzymatic reaction in glutathione synthesis driven by gamma-glutamylcysteine synthetase (γ-GCS) that has been shown to increase sensitivity to radiation.^20^ We assessed proliferation in each cell line after treatment with radiation and BSO. Radiation caused a decrease in proliferation in all 4 cell lines, whereas a glutathione depletion with a sublethal dose of BSO further sensitized only U87 and U251 to radiation (Fig1B). Radiation induced oxidative stress and double stranded DNA breaks induce cell-cycle arrest, senescence or cell death from multiple mechanisms including apoptosis, necrosis, ferroptosis and others.^19^ In the case of apoptosis, PARP (poly[ADP-ribose] polymerase) is a nuclear enzyme that detects DNA strand breaks and facilitates base excision repair by catalyzing the transfer of ADP-ribose units from NAD+ to target proteins.^21^ To assess apoptotic cell death, we probed cleaved-PARP in the cell lines radiosensitized with glutathione depletion 48 hours after treatment (Fig1C) and saw that only U251s underwent increased apoptosis in response to radiation and glutathione depletion.

**Figure 1.**
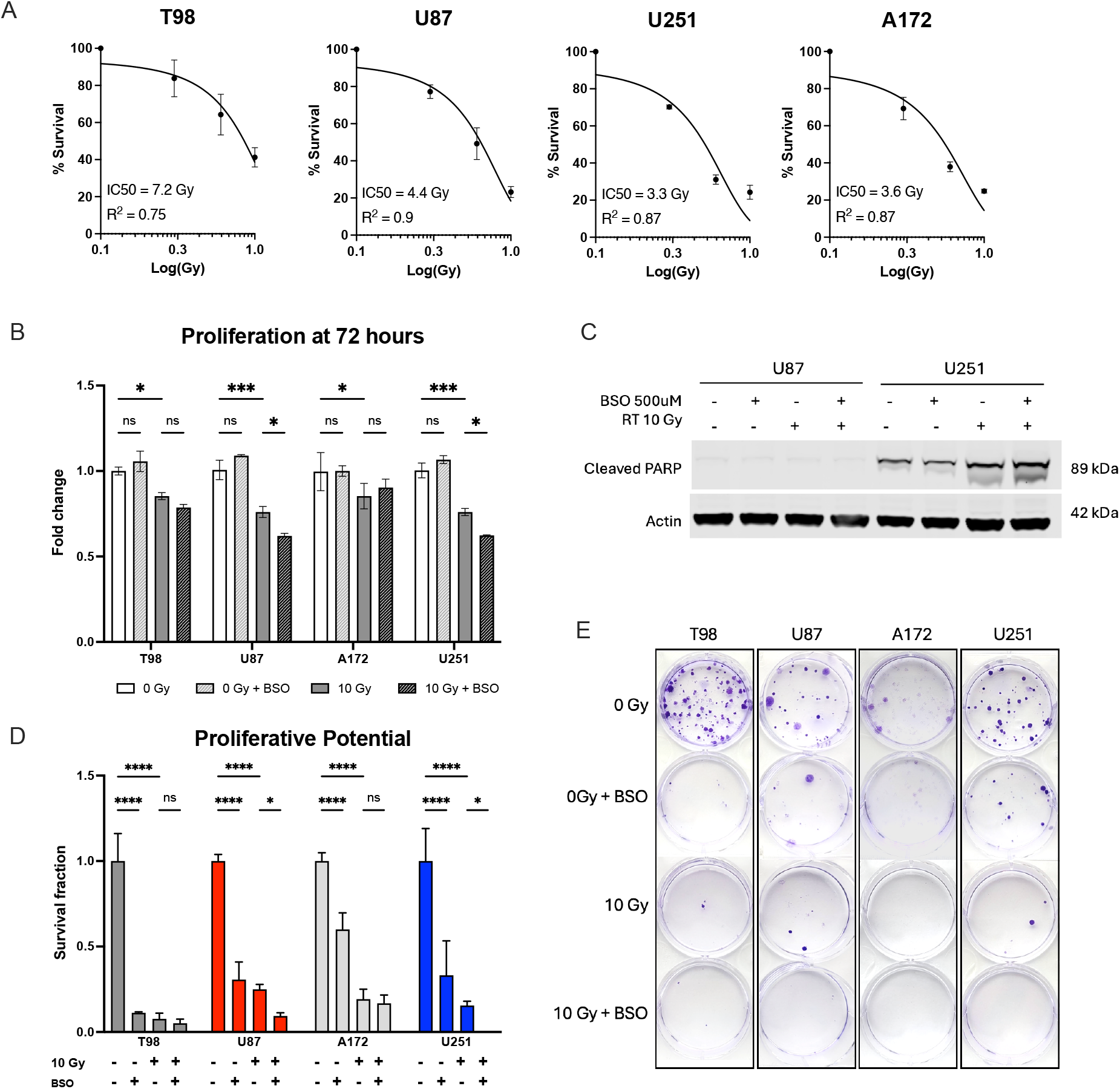
GBM radiation response in vitro. (A) Cell viability 72 hours after treatment with radiation with nonlinear fit with variable slope to obtain an estimate of IC50 and R^2^. (B) Proliferation measured by fold change in confluence after 72 hours normalized to untreated control. (C) Western Blot of cleaved-PARP in two GBM lines 48 hours after treatment with either 500 uM BSO for 18 hours, 10 Gy radiation, or both. (D) Clonogenic assays fixed and stained 2 weeks after plating, measuring proliferation potential in each condition. (E) Representative pictures of clonogenic assay wells. Note that A172 did not form as discrete of colonies as the other lines, limiting their visibility. For statistical significance in (B) and (D) 2-way ANOVA performed with significance noted by * symbols.

Proliferative potential was examined using clonogenic assays, an established *in vitro* assay for determining effects of treatment on adherent cell proliferative integrity by monitoring a single cell’s ability to proliferate. We noted all lines exhibited diminished proliferative potential after exposure to 100uM BSO for 18 hours or 10 Gy (Fig 1D). Further, the compound effect of radiation with glutathione depletion seen at 72 hours in U87 and U251 was recapitulated at 2 weeks, supporting the impact of glutathione depletion on radiation response in those GBM lines.

### B. Radiation Response in Glioblastoma is Characterized by Increase in Nucleotide Metabolism

The metabolic response to DNA damage is an upregulation of the Pentose Phosphate Pathway (PPP) to produce ribose-5-phosphate for nucleotide synthesis and NADPH which combats oxidative stress.^22^ To assess immediate metabolic changes in response to radiation, we quantified the flux through the PPP using [1,2 -^13^C] labeled glucose given immediately after radiation and measured extracellular lactate production. Glycolysis primarily produces [2,3-^13^C2] lactate ((M+2) isotopomer), while the PPP, followed by glycolysis, generates [3-^13^C] lactate ((M+1) isotopomer).^23^ We measured both the rate of PPP activity and the ratio of singlet (M+1)lactate to doublet(M+2) lactate to estimate PPP/glycolysis flux. All four GBM lines increased PPP flux (Fig 2A) and the PPP/Glycolysis ratio (Fig 2B) in response to radiation, however only U87 and U251s increased PPP flux in response to glutathione depletion. In both lines, pretreatment with BSO increased the radiation induced upregulation of PPP flux and PPP/Glycolytic flux.

**Figure 2.**
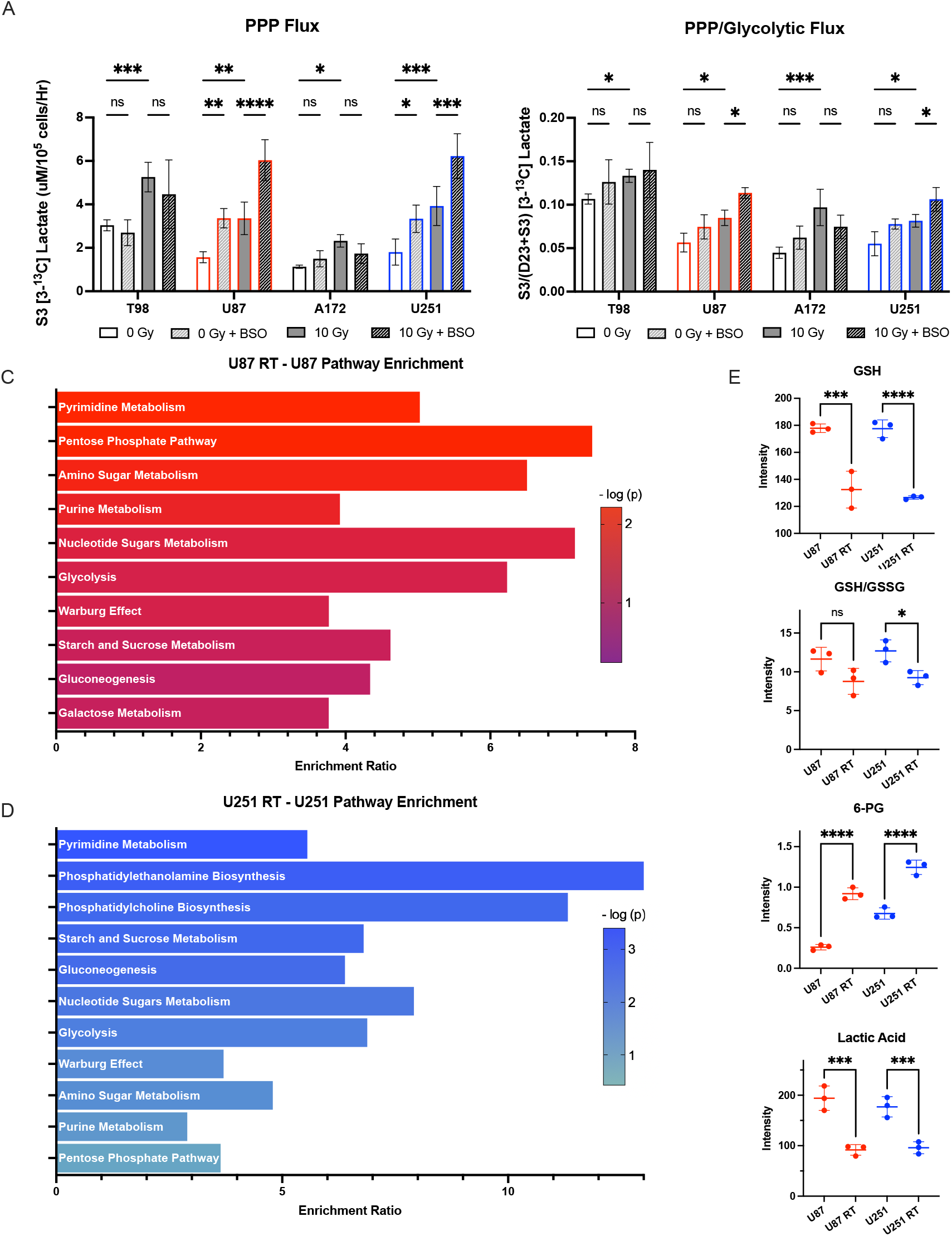
Metabolic response to radiation in U87 and U251. (A) The rate of lactate generation via the Pentose Phosphate Pathway (PPP) is plotted in each cell line treated with 500uM BSO for 18 hours, 10 Gy radiation, or both. (B) PPP/Glycolytic flux expressed as a ratio of M1/(M1+M2) isotopomers. 2-way ANOVA was performed to assess for statistical signficance. (C-D) Pathway Enrichment analyses including all pathways with p>0.1 for U87 and U251s treated with radiation compared to untreated controls of the same cell type. (E) Pool sizes from normalized intensity values of select metabolites. Significance assessed using Welch’s ANOVA test.

To further investigate the metabolic effects of radiation on glioblastoma cells in vitro, untargeted metabolomics (LC-MS) 48 hours after radiation was performed on cell extracts. Pathway overexpression enrichment analysis (Fig 2C-D) revealed U87s and U251s exposed to radiation both enriched pathways of pyrimidine metabolism (p=0.006 and p=0.0004, respectively) and purine metabolism (p=0.015 and p=0.043, respectively). In U87s, PPP was also enriched (p=0.006). As expected in response to increased oxidative stress, metabolite pool size analysis (E-H) revealed glutathione was significantly decreased in both GBM lines exposed to radiation when compared to untreated controls (E), however only U251s exhibited an imbalance in GSH/GSSG oxidative stress balance, suggesting a unique vulnerability to oxidative stress. Both lines had increased pool sizes of 6-PG, a key intermediate in oxidative PPP which supports both nucleotide synthesis and oxidative stress resistance (G). Decrements in lactate generation after radiation were seen in both GBM lines (H) and supporting prior work on this topic.^24,25^ Since the last step of anaerobic glycolysis reducing pyruvate to lactate depends on NADH, the activation of PARP and resultant depletion of NAD+ drives competition for NAD(H) availability, contributing to lower lactate as previously described in lymphoma tumors treated with etoposide.^26^

### C. Characterization of differential radioresponsivity in orthotopic xenografts in vivo

To investigate the metabolic consequences of radiation *in vivo*, we selected U251 and U87 lines for orthotopic xenograft implantation. Tumor sizes were measured twice weekly after stereotactic injection via MRI using T_2_-weighted imaging. Tumor volumes were measured in the coronal plane. Mice were treated with 0Gy (control) or 8Gy whole brain radiation in one dose 1-3 days after tumor detection on MRI. Mice were monitored for weight and neurologic morbidity and sacrificed at pre-determined thresholds for each. Tumor volumes were plotted from day of tumor detection to remove the variability with tumor engraftment time. Tumor volume curves were fit with an exponential nonlinear least-squares fit and compared using F-test for statistically significant differences. Both U87 and U251s treated with radiation demonstrated significantly slower growth rates than their untreated counterparts (*F*(2,73)= 90.19, *p*<0.0001 and *F*(2,63)= 33.22, p<0.0001 respectively) (Figure 3A-B). Growth curves for U251s treated with radiation were significantly different from U87s treated with radiation (*F*(2,78)= 376.5, p<0.0001). Radiation slows the growth of both tumor types, though much more pronounced in U251s. Further, all of the radiated U251s decreased in size in response to radiation (Fig 3B). Survival analyses demonstrated a median overall survival benefit of 9 days in U87s treated with radiation when compared to untreated controls (U87 mOS = 12 days, U87 RT mOS = 21 days) (Figure 3C) and a median overall survival benefit of 17.5 days in mice with U251s treated with radiation (U251 mOS = 14 days, U251 RT mOS = 31.5 days) (Figure 3D) when compared with untreated controls (Log Rank for all groups: X^2^=14.52, p <0.0001). Radiation conferred a longer survival benefit than in U251s than in U87s (Figure 3E), so when combined with the tumor growth rate data, U251s were deemed more radiosensitive than U87s in vivo. In a separate cohort of mice, representative brains from each group were extracted and flash frozen 48 hours after 0Gy or 8Gy radiation. A modest qualitative decrease in proliferation was seen in radiosensitive U251s treated with radiation, alongside an increase in ds-DNA damage and apoptosis (Ki-67 and γ-H2AX respectively, Figure 3G), which were not seen in radioresistant U87s. Cleaved-PARP expression was increased in both tumor types 48 hours after radiation (Fig 3F).

**Figure 3.**
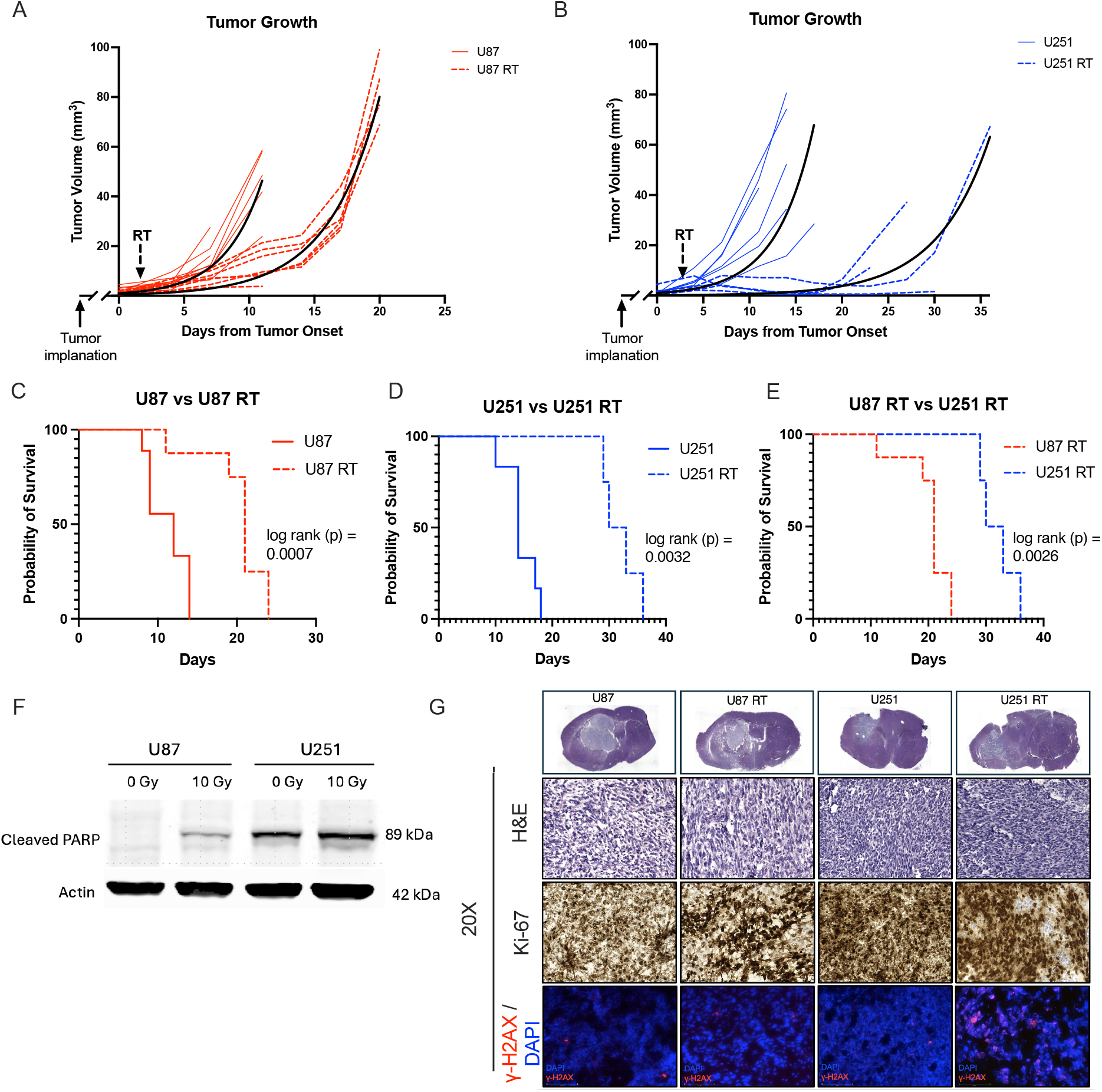
U251s are radiosenstive in vivo compared to U87. Tumor volume curves of mice with U87 (A) or U251 (B) orthotopic xenografts treated with 8Gy radiation or 0 Gy. Each line represents one mouse. Mouse numbers per group as follows: U87 = 8, U87 RT = 7, U251 = 6, U251 RT = 4. Black line is nonlinear fit for each group with Y0 constrained to <1. (C-E) Kaplan-Meier survival curves of the same mice from (A-B), a significantly increased survival benefit was seen in radiated U251s compared to radiated U87s. (F) Western Blot of cleaved-PARP from tumors 2 days after treatement with radiation. (G) Representative whole brain and 20x images of H&E, ki-67 staining, and γ-H2AX immunoflourescence in flash frozen mouse brains bearing tumors from the same treatment groups.

### D. Glioblastomas Resistant to Oxidative Stress Increase Purine Metabolism

Pyrimidine metabolism has been identified as a therapeutic vulnerability in brain tumors,^27,28^ and upregulation of purine metabolism has been linked to radioresistance in GBM which is now being studied clinically.^29,30^ To further validate our treatment-responsive and treatment-resistant models exhibit clinical relevant phenotypes, we sought to compare the metabolic consequences of radiation for each tumor. We performed unlabeled non-targeted Liquid Chromatography-Mass Spectrometry (LC-MS) analysis of mouse brain metabolome from flash frozen athymic nude mice brains with tumors (n=3 for each group). We examined relative abundance and statistical significance of radiated tumors versus their untreated controls of the same tumor type (Fig 5A-B). Radiosensitive U251s demonstrated significant increases in pyrimidine intermediate UDP and thymine in response to radiation (Fig 4B) which was expected based on our *in vitro* pathway enrichment analysis (Fig 2D). However, radioresistant U87s *in vivo* demonstrated a different phenotype and metabolic profile than *in vitro*, with a decrement in pyrimidine metabolism intermediate UDP (U87RT/U87 FC=0.46, p=0.01) and a modest but significant increase in core purine base (FC=1.16, p=0.06) (Fig 4A). Pool sizes of select metabolites further support the upregulation of pyrimidine metabolism in radiosensitive tumors and purine metabolism in radioresistant tumors (Fig 4D), reinforcing the validity of these models to recapitulate the biologic consequences of treatment response.

**Figure 4.**
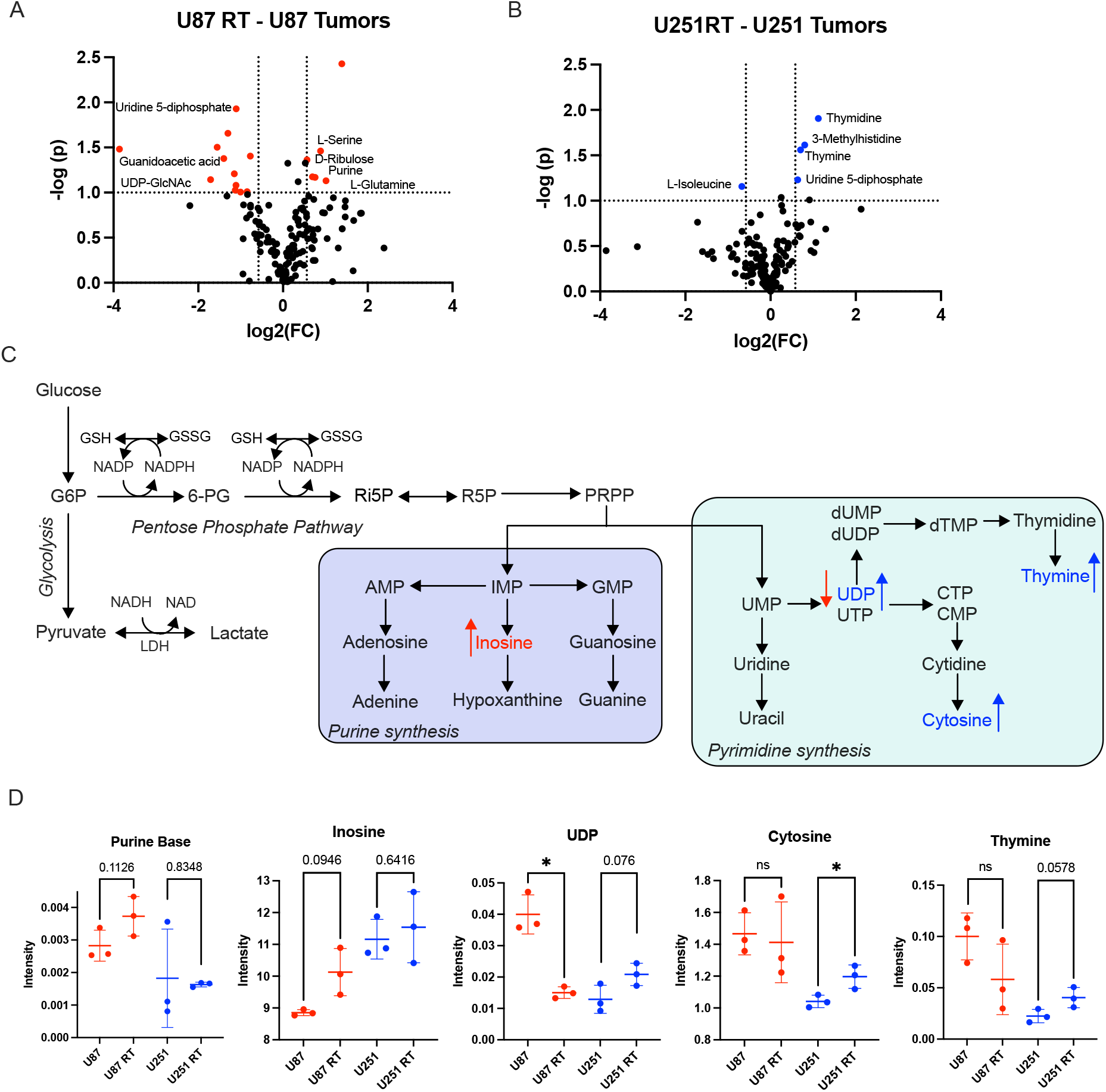
Tumors responding to radiation upregulate pyrmidine synthesis. (A-B) Volcano plots of significant metabolites from flash frozen mouse brain tissue with tumors. Thresholds of p<0.1 and FC >1.5 (<0.66) were used to identify trends towards significance. (C) Figure depicting trends in increased pyrimidines measured in radiosensitive U251s (blue) and increased purine metabolism in radioresistant U87s (red). (D) Pool sizes from normalized intensities of select metabolites. Significance assessed using unpaired t-tests between treated and untreated tumors of the same type.

**Figure 5.**
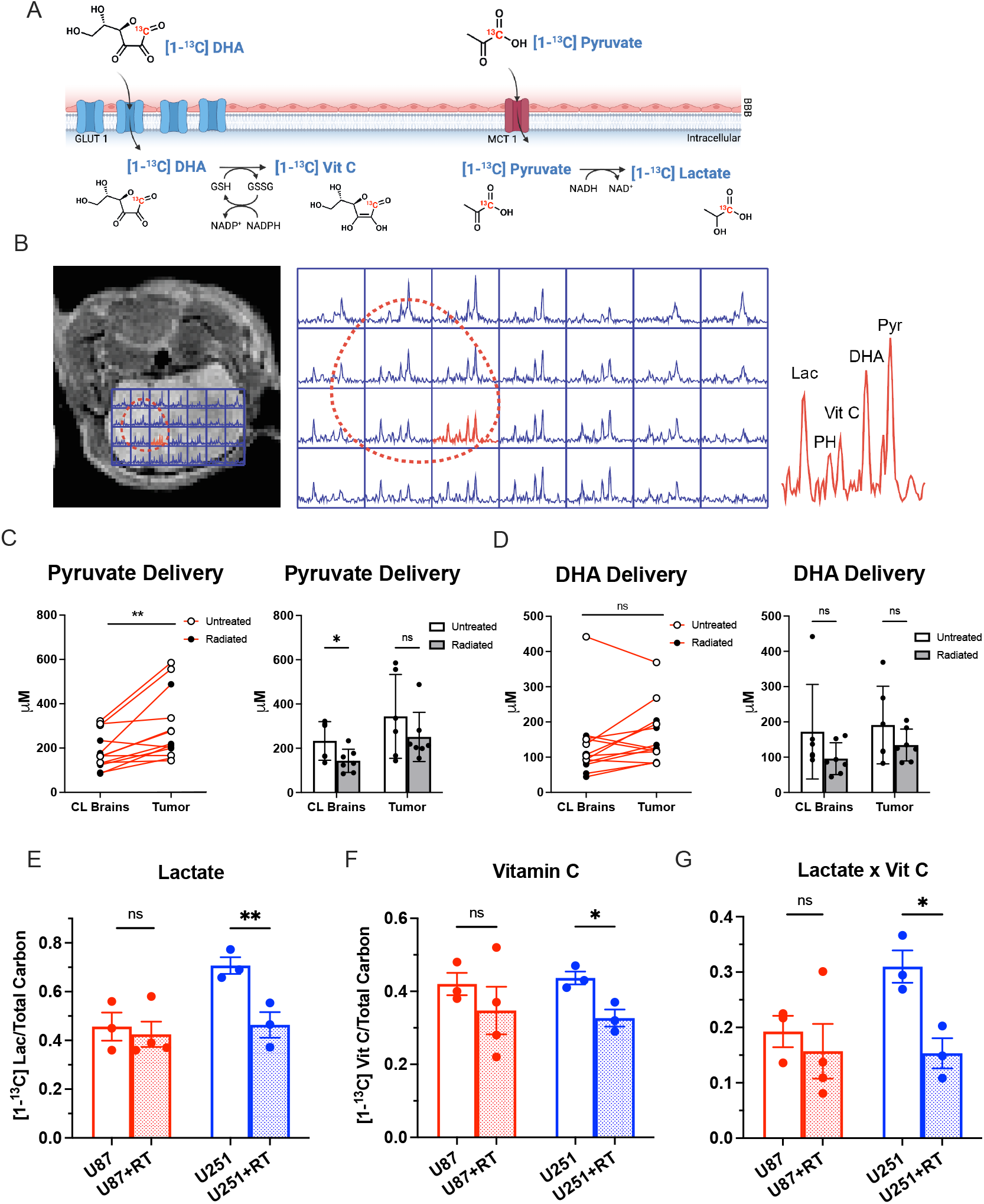
Hyperpolarized dehydroacrobate and pyruvate detect radiation response in GBM. (A) DHA’s intracellular conversion to vitamin C requires GSH and NADPH, and pyruvate to lactate requires NADH. *Made with biorender*.*com*. (B) Representative coronal T2-weighted image of U87 tumor brearing mouse brain with 2D CSI spectra of HP DHA/PA overlayed, tumor outlined in red dashed line. Representative spectra highlighted in red with products and substrates labeled. Pyr: Pyruvate. PH: Pyruvate Hydrate. Lac: Lactate. (C) Pyruvate perfusion in paired tumors and contralateral (CL) brains demonstrates differential substrate delivery, whereas DHA perfusion is unchanged (D). Lactate (E) and Vitamin C (F) generation from 4 cohorts of mice (n=3 each except U87 RT which had n=4 mice) demonstrating a decrement in both Vit C and lactate in radioresponsive tumors. Lactate * Vit C (G) may serve as a single biomarker to predict treatment effect in GBM. Signficance in (E-G) assessed using unpaired t-tests comparing treated to untreated groups of the same tumor type.

### E. Decrements in Reductive Capacity and Glycolytic Flux Predict Response to Oxidative Stress in Glioblastoma

Based on the demonstrated sensitivity to glutathione availability and decrement in lactate production in the response to radiation, we hypothesized that probing reductive capabilities of tumors would reveal early metabolic effects of radiation. To accomplish this, we utilized co-hyperpolarized [1-^13^C] dehydroascorbic acid and [1-^13^C] pyruvic acid. Dehydroascorbic acid (DHA) is readily transported across the blood-brain barrier and enters the cell through GLUT1 where it is rapidly two-electron reduced to Vitamin C (Vit C) in a glutathione and NADPH dependent reaction (Fig 5A). HP [1-^13^C] DHA, therefore, can be used to assess the capacity of tumors to resist oxidative stress. HP DHA has demonstrated ability to act as a redox sensor in multiple preclinical models^8,9,11^ including prostate cancer.^7^ We have developed co-polarized DHA with pyruvic acid, the most widely used HP imaging probe. HP Pyruvate enables real time assessment of glycolytic flux via quantification of lactate generation (among other products), which has been studied clinically in tumor diagnosis, predicting tumor grade, and treatment response in brain, breast, renal, and prostate cancer. ^31-34^ Combining HP DHA with HP PA, unique chemical shifts allow us to spatially resolve both substrates and their respective products (lactate and Vitamin C) using spectroscopic imaging (Fig 5B).

To examine regional variations in tissue perfusion, a distinct limitation of existing clinically used methods, we compared the spatial differences brain tumors compared to the contralateral (CL) brain. Using methods we have previously published,^13^ absolute concentrations of each product were calculated. Pyruvate delivery was higher in tumors and in untreated compared to treated tumor or CL brain regions (Fig 5C). This differential substrate delivery is a key limitation of the clinical interpretation of HP pyruvate readouts. However, DHA substrate perfusion was unchanged between tumors and CL brain, nor was it affected by radiation (Fig 5D), highlighting the advantage of adding this probe with existing methods. In untreated mice, there was no difference in substrate perfusion between either tumor type compared to CL brain (Supplementary 1), highlighting the role of radiation in substrate delivery.

Treatment-sensitive U251 tumors treated with radiation led to decrement in the conversion of DHA to Vit C and in pyruvate conversion to lactate when compared to untreated controls, reflective of diminished reductive capacity of these tumors (Fig 5C). Correspondingly, treatment-resistant U87 tumors generated no difference in relative lactate or Vit C generation after radiation, reflecting maintenance of their reductive capacity in the face of oxidative stress. We investigated the product of Lactate/Total Carbon and Vit C /(DHA + Vit C) as a combined readout to capture the activity of both pathways simultaneously, and found it was likewise significantly lower in radiosensitive U251 tumors, highlighting its potential use as a single biomarker for treatment response (Fig 5D).

## II. Discussion

We present the first study utilizing HP DHA as a redox sensor in brain tumors and demonstrate the quantitative measure of oxidative stress using HP DHA/PA as a tool to identify treatment response in glioblastoma that overcomes limitations of existing clinical methods. We validate the chosen models by demonstrating the responses to oxidative stress *in vitro* and *in vivo* align with known mechanisms of response to radiation, which supports their use in biologic validation of an oxidative stress imaging probe.

The metabolic response to radiation is characterized by increase in PPP flux, generating both nucleotide precursors to repair DNA damage and NADPH as a reducing agent to combat intracellular oxidative stress.^35^ Our *in vitro* data are concordant with this paradigm. Cellular response to radiation in these four GBM lines was characterized by increase in PPP rate and relative flux when compared to glycolysis explaining the decrement in lactate generation seen in pool size analysis. NADPH is a product of driving PPP forward, supporting the need for antioxidants in response to radiation induced oxidative stress. Along with NADPH and NADH, reduced glutathione (GSH) is another key antioxidant that maintains cellular redox homeostasis. Decreased glutathione pool sizes seen in both radiated cell lines is consistent with the consumption of reducing agents in response to oxidative stress. Further, we demonstrate that glioblastoma sensitivity to radiation is augmented by glutathione depletion, reinforcing the potential utility of a redox sensor to measure radiation response.

Phenotypic differences in the response to radiation in U87s emphasize the importance of *in vivo* studies in this group. Physiologic variables associated with *in vivo* tumors including the tumor microenvironment, perfusion, impact of immunogenicity, methodologic techniques such as the time to extraction, and low sample sizes likely contributed to the higher variance seen in tissue metabolomics data compared to *in vitro*. More liberal significance cutoffs for p-value (>0.1) and fold change (<0.5 and >1.5) were used to elucidate metabolic trends, though the generalizability of these findings remains to be seen. Despite this variability, trends in metabolic phenotypes still complement the existing literature and support their utility as suitable models for imaging probe validation.

Inhibition of de novo pyrimidine synthesis has been investigated clinically in cancers with disappointing results.^36^ However, a newly targeted intermediate in this pathway has been recently exploited as a therapeutic vulnerability in brain cancers^27,28^ by targeting dihydroorotate dehydrogenase (DHODH), a key intermediate in the synthesis of uridine monophosphate (UMP), a precursor for thymidine and cytosine. Shi et al. demonstrated IDH-mutant (IDHmt) tumors were more vulnerable to the imbalance of pyrimidines and purines induced by DHODH inhibition than IDH-wild type,^37^ and Pal et. al. showed that adult-type glioblastoma was not as vulnerable to DHODH inhibition as diffuse midline glioma.^38^ Together, these studies suggest that GBMs, which are clinically inherently resistant to treatment, may be less dependent on pyrimidine synthesis upregulation. In line with this, our data demonstrated that only relatively treatment-sensitive adult-type glioblastoma U251s (IDHwt) upregulated pyrimidine synthesis in response to radiation.

In contrast to the radiosensitive U251s, the relatively radioresistant U87s favored purine metabolism with a significant decrement in UDP and increase in purine bases and inosine. This aligns with known mechanisms of radioresistance in glioblastoma. For example, in a metabolomic study of 23 glioblastoma cell lines, Zhou et. al. correlated increases in purine metabolism with radioresistance and demonstrated the inhibition inosine monophosphate dehydrogenase (IMPDH1) specifically sensitizes radioresistant GBM models to radiation, including U87s.^29^ They went on to correlate lower IMPDH1 gene expression with poorer prognosis in The Cancer Genome Atlas Database, a finding which was validated in two other large international cancer genomic databases.^39^ Our work strengthens the support for metabolic dependence on purine metabolism in treatment-resistant glioblastoma and reinforces the use of radiation treatment in U87s as a model resistant to oxidative stress.

Hyperpolarized [1-^13^C] pyruvate is being clinically investigated as a tool to predict treatment response in a variety of cancer indications. The basis of these investigations is founded in the Warburg effect, in which cancer cells preferentially convert pyruvate to lactate even in the presence of functioning mitochondria and oxidative phosphorylation. Thus, quantifying this reaction has been shown to predict tumor grade, distinguish tumor from nontumor tissue, and identify treatment responders.^31-33,40^ The NADH dependent conversion of pyruvate to lactate catalyzed by LDH serves as a marker both for glycolytic flux and an indirect quantification of NADH availability. We utilize co-polarization with [1-^13^C] dehydroascorbate, which undergoes rapid NADPH and glutathione dependent reduction to Vitamin C. Thus, the availability of reducing agents (GSH, NADPH, and NADH) can be detected by quantifying the conversion of pyruvate to lactate and DHA to Vit C. Given known induction of oxidative stress by ionizing radiation, the influence of glutathione availability on radiation response *in vitro*, and decrement in lactate production after treatment, we identified HP DHA/PA as an ideal imaging biomarker to quantify the response to radiation and identify treatment responders before anatomic changes could be visualized.

2D-Chemical Shift Imaging after injection with HP DHA/PA revealed a decrement in lactate and Vitamin C production in radiosensitive U251. This is contrary to findings in a previous study utilizing hyperpolarized pyruvate which did not detect a difference in lactate production in orthotopic models of U251s treated with radiation,^41^ which may be due to a lower dose of radiation (6Gy compared to 8Gy in our study) or to evaluation at an earlier time point. Mair et. al. quantified lactate generation using HP pyruvate at baseline and in response to radiation in U87 tumors along with four patient derived xenografts, revealing correlation of lactate labeling with c-Myc expression and glycolytic enzyme expression (LDHA, HK2) which were associated with radioresistance of cell lines in vitro.^42^ A trend toward increased lactate generation at baseline in radioresistant U87s when compared to contralateral brain in our study is consistent with this framework, and maintenance of high generation of lactate after radiation suggests resistance to treatment. Likewise, a decrement in lactate production is consistent with tumors that are responding to treatment, as was seen in radiosensitive U251s in our study. Our work, taken together with existing literature, support the utility of longitudinal monitoring of metabolic profiles before and after treatment to monitor the tumor metabolic response to treatment over time.

Given the role of oxidative stress in a plethora of human diseases, the development of a non-toxic, spatially resolved and quantitative oxidative stress imaging biomarker is an unmet clinical need that would greatly improve our ability to tailor patient treatment plans. Methods to image oxidative stress *in vivo* include fluorescent probes which are difficult to implement *in vivo* and paramagnetic MRI contrast agents which are often limited by short relaxation times.^43^ Radiolabeling of fluorescent probes and imaging of downstream consequences of oxidative stress have been utilized in PET but come at the cost of radiation exposure. Prior work by our group and others utilizing HP DHA have demonstrated utility of this endogenous substrate as a redox sensor in models of a variety of malignancies including lymphoma,^14^ colorectal cancer,^15^ and prostate cancer^7^ along with nonmalignant disease states.^8,9,12^ Recently, co-polarized HP DHA/PA was shown to achieve high enough spatial resolution to distinguish distinct regional metabolic profiles between white matter and deep gray nuclei in the mouse brain.^13^

Our data show that in a state of oxidative stress, both lactate and Vit C generation will decrease in response to a bolus of their substrates. Combining these readouts provides a singular variable reflective of the overall reductive capacity of the tumor and could be used as a biomarker for treatment response. We therefore propose examining the product of lactate/total carbon and Vit C / (DHA + Vit C) as a unified parameter reflective of the overall state of oxidative stress. It is possible that there are conditions in which only one of Vit C or lactate goes down, or both decrease but only modestly. Combining these variables is expected to be more sensitive than considering each product individually and would reflect the overall reductive capacity of the tumor, simplifying the interpretation of the results. Future studies with co-polarized substrates could employ similar tactics, strengthening the clinical translatability of new HP imaging probes.

We present the first study successfully quantifying oxidative stress in glioblastoma using hyperpolarized MRI and address a key clinical limitation of existing metabolic imaging methods by optimizing substrate perfusion. Further, we identify a singular biomarker, Lactate X Vitamin C, with the ability to predict treatment response before anatomic changes can be visualized, holding potential to meet a significant unmet clinical need. Beyond applications in radiation response, many other cancer-directed therapeutics are aimed at targeting oxidative stress in brain tumors,^4^ broadening the potential applications of HP DHA both in this populations and patients with other malignancies. Moreover, with the exciting developments in targeted radioligand therapy,^2^ HP DHA/PA, holds potential to transform our ability to monitor treatment response in brain tumors in real time, and these results motivate its’ urgent clinical translation. Furthermore, early measurements of treatment response with HP DHA/PA can be used in future studies of novel treatments to capture the breadth of patient tumor responses and expedite the selection of promising treatment candidates in both preclinical and clinical spaces.

## V. Acknowledgements

We acknowledge all Keshari lab members for help with experiments or preparation of this manuscript. We thank the MSKCC cores for their help in designing and conducting experiments: Antitumor Assessment core, Molecular Cytology core, NMR Analytical core and the Animal Imaging core. Mass spectrometric analyses were performed at the Weill Cornell Medicine Proteomics and Metabolomics Core Facility. This work was supported by the National Institute of Health – P30 CA008748, R01 CA252037 (K.R.K), The Paul Calabresi Career Development Award for Clinical Oncology: K12 CA184746, and the MSK Imaging and Radiation Sciences Program (IMRAS) Seed Grant.

## Contributions

Conceptualization, K.R.K., I.K.M; methodology, K.R.K., E.C., P.P., S. P.; investigation E.C., P.P., S. P. G.F., M. B.; formal analysis, E.C., K.R.K.; writing-original draft: E.C.; writing—review and editing, E.C., P.P., S. P.; supervision, K.R.K., I.K.M; funding acquisition K.R.K. and E.C.

## Data Availability Statement

The metabolomic data generated in this study are available within the article and its supplementary data files. All other data were generated by the authors and available on reasonable request.

