## Supplemental 1 for "Co-Hyperpolarized Dehydroascorbate and Pyruvate MRI Predicts Treatment Response in Glioblastoma"

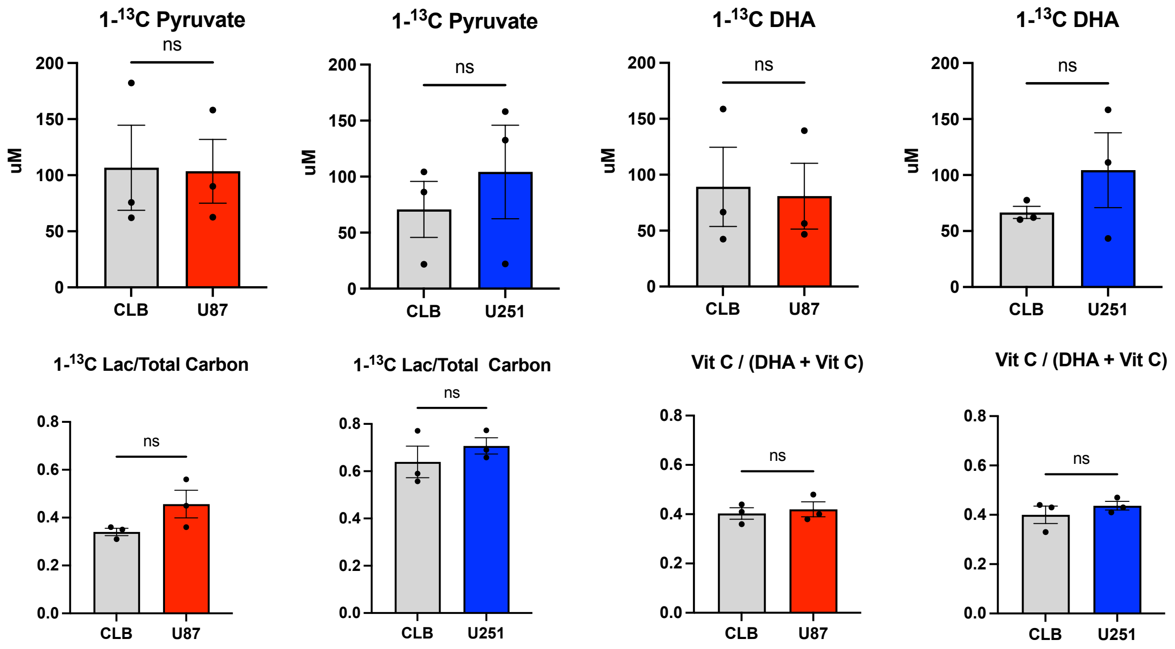


**Supplementary Figure 1:** Concentrations of substrates and ratios of products over the total carbon for their substrate detected in tumor bearing mice with U251 or U87s tumors compared to the contralateral brain (CLB) in the same mice. n=3 of each group. Students paired t-test used for statistical significance.
