## Supplemental 4 for "Co-Hyperpolarized Dehydroascorbate and Pyruvate MRI Predicts Treatment Response in Glioblastoma"

Supplemental 4: Additional methods

**Western blot:** Cells were lysed with RIPA buffer (Thermo Fisher Cat #89900) with phosphatase/protease cocktail (Thermo Fisher Cat # 78440). Lysates were quantified using DC protein assay according to manufacturer’s protocol and resolved on 4–12% Bis-Tris gels (Thermo Fisher # NP0321BOX) under denaturing conditions. Proteins were blotted onto a PVDF membrane (Thermo Fisher # 88520) and probed with anti-cleaved-PARP antibody (CST # 9544) and anti-Actin (CST # 8457S) Membranes were visualized using anti-rabbit HRP-conjugated secondary antibodies. Signals were visualized using HRP substrate. For flash frozen tumor tissues, samples underwent tissue homogenization using bead beating and sonication according to manufacturer’s instructions before undergoing identical protocols to above.

**Hyperpolarization methods:** One hundred microliters of [1-^13^C]DHA/[1-^13^C]PA (40/60; v/v) doped with 15 mM AH111501 trityl radical was prepared. The sample was polarized for 1.5 to 2 hours on a SpinLab hyperpolarizer (5 T, 0.8 K, GE HealthCare) by microwave irradiation. After polarization, the frozen sample was dissolved D_2_0 and pyruvate was neutralized by adding sodium acetate solution (in D_2_O) to the dissolute to afford the HP solution of pH ~ 5.5. The HP solution contained 40 mM [1-^13^C]DHA and 90 mM [1-^13^C]PA. Approximately 700 μl of the HP solution was transferred to a 5-mm NMR tube and loaded to a 1-T Spinsolve ^13^C NMR spectrometer (Magritek, Wellington, NZ). The spin-lattice relaxation time (T1) was estimated using a monoexponential fit and corrected for the flip angle. Thermal polarization was determined from the average spectrum of 256 scans acquired with a flip angle of 90° every 10 s. Final polarization values, corrected for flip angles and the apparent T1, averaged 22% for pyruvate and 17% for DHA. The concentration of ^13^C-DHA and ^13^C-PA in the HP solution was measured by ^13^C NMR at 600 MHz in the presence of mM Gd-DOTA and [1-^13^C]lactate standard.
